# Is diversity in worker body size important for the performance of bumble bee colonies?

**DOI:** 10.1101/2020.05.06.079525

**Authors:** Jacob G. Holland, Shinnosuke Nakayama, Maurizio Porfiri, Oded Nov, Guy Bloch

## Abstract

Specialization and plasticity are important for many forms of collective behavior, but the interplay between these factors is little understood. In insect societies, workers are often predisposed to specialize in different tasks, sometimes with morphological or physiological adaptations, facilitating a division of labor. Workers may also plastically switch between tasks or vary their effort. The degree to which predisposed specialization limits plasticity is not clear and has not been systematically tested in ecologically relevant contexts. We addressed this question in 20 freely-foraging bumble bee (*Bombus terrestris*) colonies by continually manipulating colonies to contain either a typically diverse or reduced (“homogeneous”) worker body size distribution, over two trials. Pooling both trials, diverse colonies did better in several indices of colony performance. The importance of body size was further demonstrated by the finding that foragers were larger than nurses even in homogeneous colonies with a very narrow body size range. However, the overall effect of size diversity stemmed mostly from one trial. In the other trial, homogeneous and diverse colonies showed comparable performance. By comparing behavioral profiles based on several thousand observations, we found evidence that workers in homogeneous colonies in this trial rescued colony performance by plastically increasing behavioral specialization and/or individual effort, compared to same-sized individuals in diverse colonies. Our results are consistent with a benefit to colonies of predisposed (size-diverse) specialists under certain conditions, but also suggest that plasticity or effort, can compensate for reduced (size-related) specialization. Thus, we suggest that an intricate interplay between specialization and plasticity is functionally adaptive in bumble bee colonies.

## INTRODUCTION

A key organisational principle of insect societies is division of labor, whereby individuals specialize in various tasks such as caring for the brood (“nursing”), guarding the nest, or foraging for resources, and disproportionately perform these activities compared to the group average (Beshers and Fewell, 2001; Wilson 1971). Division of labor is thought to improve colony-level efficiency if the costs of task-switching are high, or if generalists perform tasks less efficiently than specialists (Anderson and McShea, 2001; Goldsby et al., 2012; Oster and Wilson, 1978; West et al., 2015). Similar benefits are also found in other collective systems, including non-insect animals (e.g. Arnold et al., 2005; Bennett and Faulkes, 2000), cells within multicellular organisms (Maynard Smith and Szathmáry, 1995), cells within bacterial colonies (Zhang et al., 2016), and human societies (Nakayama et al., 2018; Smith, 1776). Social insects provide outstanding model systems to unravel the mechanisms and organisational principles of division of labor because colony performance has been shaped by strong selection and can be manipulated in ecologically-relevant contexts.

In many insect societies, division of labor results from individuals with inherent behavioral specialization, based on developmentally or genetically determined variation, which we refer to as “predisposed specialists”. Predisposed specialization may be accompanied by morphological, anatomical, or physiological features suited to particular behavioral roles. Extreme examples include size differences of more than fiftyfold in the morphological castes of ant workers (Ferguson-Gow et al., 2014) and developmental morphological changes in many termites (Korb and Hartfelder, 2008), as well as physiological changes associated with age in adult workers of many social hymenopterans (Robinson, 1992; Wilson 1971). These individual differences are assumed to be functionally linked to specialization, allowing workers to be more efficient at performing certain tasks, but may come at the expense of decreasing individual behavioral plasticity (e.g. Johnson, 2003; Moglich and Holldobler, 1974; Robinson et al., 2009; Tofts and Franks, 1992). In particular, it could create a need to produce balanced proportions of different predisposed specialists within a colony, so that all tasks are performed in accordance with colony requirements. Presumably, this is why such predisposed specialists are normally associated with large colonies, where there are enough individuals to provide a buffer against unexpected changes (Anderson and McShea, 2001; Bourke, 1999; Ferguson-Gow et al., 2014).

In smaller colonies with predisposed specialists, rapid changes in colony demands or disproportionate mortality of certain specialized workers could lead to inappropriate ratios of specialists. This may mean that retaining at least some level of plasticity is more crucial in small societies. Behavioral plasticity has indeed been shown to play an important role in several social systems, and often appears to be compatible with a response-threshold model (Beshers and Fewell, 2001; Johnson, 2003; Robinson et al., 2009). In simple form, this describes nestmates differing in internal sensory thresholds to task-related stimuli, meaning that moderately sensitive individuals will perform a task in the absence of highly sensitive individuals, via an increase in the appropriate stimulus (Beshers and Fewell, 2001; Page et al., 1998; Weidenmuller, 2004).

It is assumed that some forms of specialization such as large or small body size can limit the ability to switch tasks, but the degree to which predisposed specialization limits plasticity is not clear (Johnson, 2003). Furthermore, the interplay between morphological specialization and plasticity has not been systematically tested in ecologically relevant contexts (Jeanson and Weidenmuller, 2014). We addressed this issue using colonies of the bumble bee *Bombus terrestris*, which are especially well suited for studies on colony performance because of their relatively small size, annual cycle, and amenability to precise social manipulations in an ecologically relevant context. For example, nests can be kept in controlled conditions to support basic colony needs, while workers freely-forage in the natural environment (e.g. Lopez-Vaamonde et al. 2004; Gill et al., 2012; Shpigler et al., 2016; Yerushalmi et al., 2006). Their social biology also lends itself to testing this relationship, because there is good evidence for predisposed specialization based on body size, but there is also reason to expect individual plasticity. In particular, there is typically a behavioral continuum from the smallest to the largest workers, which tend towards specialization in nursing and foraging, respectively (Alford, 1975; Brian, 1952; Cumber, 1949; Free, 1955; Gardner et al., 2007; Goulson et al., 2002; Yerushalmi et al., 2006; for a recent review see Chole et al. 2019, in press). Large body size is also associated with anatomical and morphological features which appear to contribute to increased foraging performance (Klein et al., 2017; Spaethe and Weidenmuller, 2002). For example, large workers possess larger eyes (Kapustjanskij et al., 2007; Spaethe and Chittka, 2003), larger brains (Mares et al., 2005; Riveros and Gronenberg, 2010) and a greater density of antennal sensilla (Spaethe et al., 2007) which are associated with better visual and olfactory acuity. They also have more cells expressing the circadian neuropeptide Pigment Dispersing Factor (PDF; Weiss et al., 2009) and, concordantly, stronger diurnal circadian rhythms (Yerushalmi et al., 2006). There is also evidence suggesting that smaller bees may be better suited to performing some in-nest activities (Couvillon and Dornhaus, 2010). This individual variation is largely independent of genetic effects, since bumble bee colonies are typically headed by lone singly-mated queens and so workers are closely related. Furthermore, ultimate body size and larval developmental duration are determined largely by the social environment in which the brood develops and not by its source colony or factors in the egg (Shpigler et al., 2013). Overall, this predisposed behavioral specialization between workers, with the associated size and morphological variation, could limit individual plasticity. However, bumble bee colonies are relatively small, typically growing from ten or fewer workers in the first brood to no larger than a few hundred workers at peak size (Alford, 1975; Goulson, 2010). Thus, plasticity in worker behavior could be important for allowing bumble bee colonies to appropriately respond to changes in the environment or in colony composition. Consistent with this prediction, individual workers have been long known to perform a variety of different tasks (e.g. Alford 1975), and reducing the numbers of specialist workers did not reduce thermoregulation and undertaking performance in laboratory-confined colonies of *Bombus impatiens* (Jandt and Dornhaus, 2014).

Manipulating colony composition can be a useful approach for testing the functional significance of size-based specialization in social insects (Wills et al., 2018), and has been used successfully in field studies with ants (e.g. Billick, 2002) and in laboratory settings with both ants and bees (e.g. Billick and Carter, 2007; Jandt and Dornhaus, 2014). Here, we created colonies with a diverse or homogeneous body-size distribution of workers and monitored them in freely-foraging conditions to test the following hypotheses. Hypothesis 1: *Wide size variation, associated with predisposed specialization, improves colony performance because morphologically diverse individuals are adapted to the tasks in which they specialize*. Thus, “homogeneous” colonies, in which the size-diversity of workers is experimentally reduced, are predicted to perform worse than typically size-diverse colonies. Hypothesis 2: *Homogeneous colonies attempt to compensate for the loss of predisposed specialists.* We predicted that this would be achieved by the following non-mutually exclusive mechanisms: a) increasing the production of predisposed specialists (i.e., small or large individuals); b) increasing the proportion of specialists via behavioral plasticity; and/ or c) increasing effort in each individual (e.g., reducing inactivity or increasing foraging effort).

## METHODS

### Colony Maintenance and Treatment

Incipient *B. terrestris* colonies were purchased from Pollination Services Yad Mordechai, Kibbutz Yad Mordechai, Israel. Trials 1 and 2 used different cohorts of colonies and began on 14 May 2015 and 26 June 2016 (each designated ‘Day 1’), respectively. In Trial 1, the nine focal colonies contained a queen and 2-6 workers, and in Trial 2, the eleven focal colonies contained a queen and 1-7 workers. The trials were broadly similar, although several differences existed; for example, the colonies in Trial 2, but not Trial 1, were closely genetically related to each other. Further details of this and other aspects of colony rearing, are summarised in the Supplementary Methods and in Table S1.

All colonies were housed in large wooden boxes from day 2 of the experiment in one of three environmental chambers in the Bee Research Facility of the Edmond J. Safra campus of the Hebrew University of Jerusalem, Givat Ram, Jerusalem. The chambers were maintained at approximately 28°C (mean ± SD; Trial 1 = 27.7 ± 0.7 °C; Trial 2 = 28.6 ± 0.9 °C) and 50% relative humidity (RH; mean ± SD; Trial 1 = 49.0 ± 6.7%; Trial 2 = 47.3 ± 11.6%) for the duration of experiment. Within these chambers, nests were maintained in constant darkness or dim red light for the majority of the time, with dim white light used for behavioral observations. On day 12 or 13 (Trial 1) or day 17 (Trial 2) of the experiment, each colony was connected to the outside environment via plastic tubing (approximately 120 cm long), passing through the external wall of the environmental chamber and ending in a landing pad, which allowed the bees to freely forage in the campus and surrounding area. The colonies were located in close proximity to each other and to our honeybee apiary, which is known to be a challenging environment for bumble bees due to competition over resources (Elbgami et al., 2014). The hot dry climate of the Jerusalem summer was also expected to be challenging for *B. terrestris*, which has a largely temperate distribution. Such challenging conditions enabled us to clearly assess the influence of treatment on colony fitness.

Colonies were initially provided with ad libitum inverted sugar syrup and pollen (mixed to a thick paste with a small amount of syrup), provided by the colony supplier. After being connected to the outside, colonies were given decreasing amounts of supplementary pollen and syrup, equally across colonies. In Trial 1, provisioning with pollen and syrup was stopped at Day 22 and 24 respectively (i.e. 10 and 12 days after the first colonies were connected). In Trial 2, which was conducted later into the summer, a small amount of pollen and syrup continued to be provided, equally across colonies, throughout the duration of the experiment. Specifically, 0.5-1 g of pollen and/or 1-5 ml of syrup was provided directly into food cells of each colony every 1-5 days, depending on stored food levels, with equal amounts always supplied to all colonies on any given day. This supplementary food (Trial 2 only) was provided to prevent the colonies from collapsing, since almost all colonies were struggling to grow (i.e. only a slight or no increase in total workers beginning shortly after the colonies were connected). The amount provided was not sufficient to support the colonies completely (indeed all colonies remained fairly weak) and, for most colonies across both treatments, was only a small proportion of the total food they collected. Thus, all colonies still needed to forage for food, and it is unlikely that this supplementary food in Trial 2 would compensate for any effect of treatment on colony performance. Until the time of colony connections, any males or dead workers found in colonies were replaced by similarly sized workers as part of worker redistribution the following day (see Figure 1 and below), in order to mitigate any effect of male production or mortality at this stage on colony performance.

**Figure 1.**
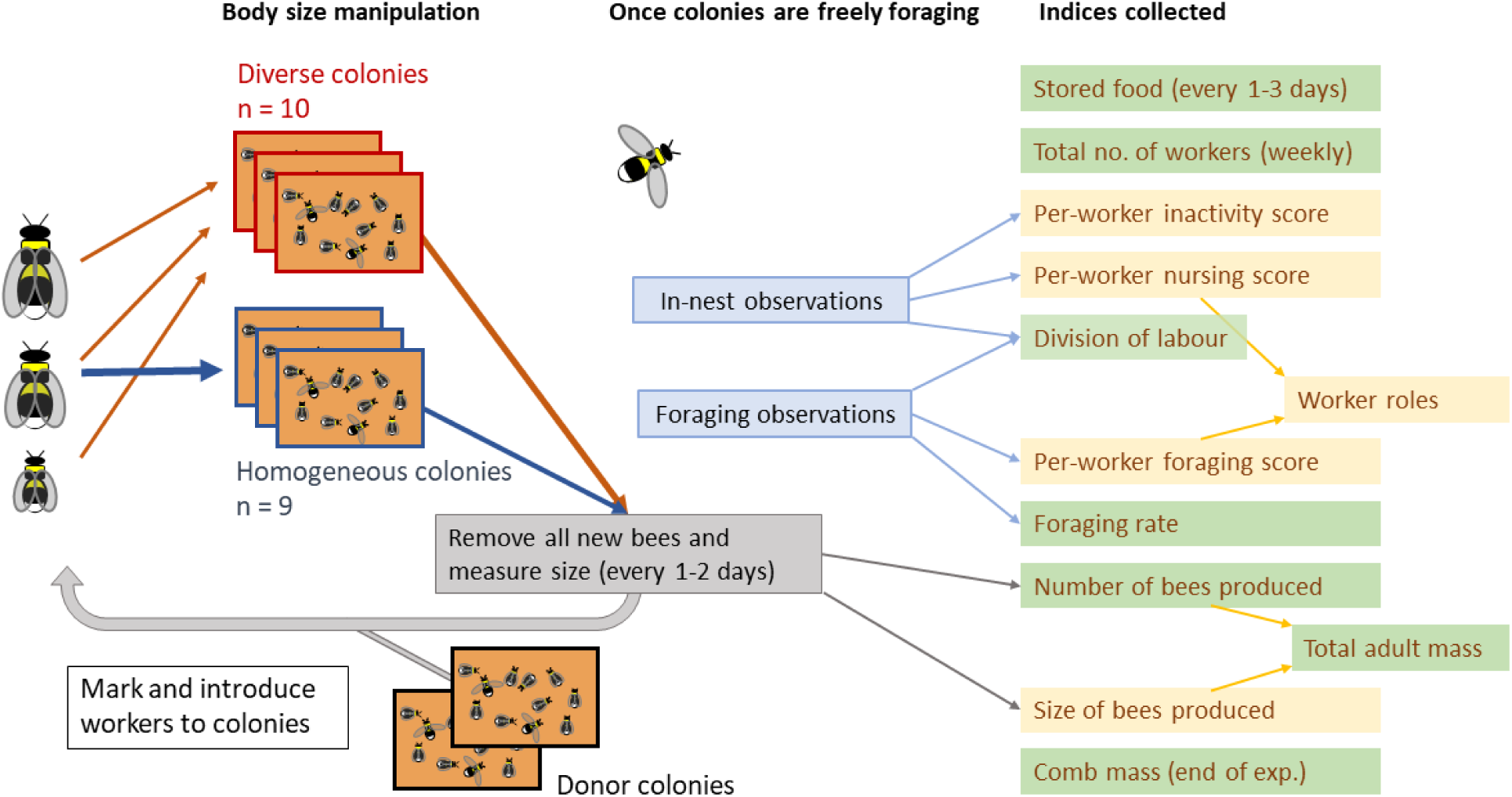
Experimental design summary. Lab colonies of *Bombus terrestris* were continuously manipulated over about two months, with all new workers redistributed according to body size to maintain either diverse (all sizes) or homogeneous (only medium size) distributions. After an initial establishment period, colonies were allowed to freely forage in the natural environment. Behavioural observations and other assessments were used to produce a number of individual-level indices (orange boxes) and colony-level indices (green boxes) which were used for assessing behaviour and colony performance in downstream analyses.

In both trials, colonies were randomly assigned to one of two treatments on Day 1: diverse or homogeneous. The initial number of workers did not differ between treatments (Wilcoxon rank sum tests: Trial 1, *W =* 15.5, *p =* 0.2; Trial 2, *W =* 12, *p =* 0.6). These treatments were created and maintained by continually collecting and redistributing workers across colonies (Figure 1; total introduced workers: Trial 1, *n =* 1149; Trial 2, *n =* 2081). Starting with the initial workers, every 1-2 days throughout the experiment, all previously unmeasured workers from all focal colonies were collected, measured and classed as small (marginal cell length ≤ 2.3 mm), medium (2.4 – 2.7 mm) or large (≥ 2.8 mm). Most of these workers were then marked and redistributed among the focal colonies in accordance with colony treatment, along with a smaller number of workers from non-focal donor colonies (Trial 1, 9%; Trial 2, 19% of all introduced workers were from genetically unrelated donor colonies). Specifically, homogeneous colonies received only medium workers, whilst diverse colonies received small, medium and large workers in similar proportions. In this way, the overall mean sizes of introduced workers was kept the same between treatments, but the standard deviations differed (mean ± SD marginal cell length of workers introduced over whole experiment: Trial 1, homogeneous colonies = 2.59 ± 0.14 mm (*n =* 622), diverse colonies = 2.59 ± 0.37 mm (*n =* 527); Trial 2, homogeneous colonies = 2.55 ± 0.10 mm (*n =* 948), diverse colonies = 2.55 ± 0.33 mm (*n =* 1133)). Furthermore, the mean size of introduced workers on any given day was also kept approximately the same between homogeneous and diverse colonies (mean ± SD daily difference between mean marginal cell lengths of workers introduced to the two treatments; Trial 1 = 0.063 ± 0.06 mm; Trial 2 = 0.021 ± 0.024 mm).

On each day, colonies in each treatment received the same number of workers, and thus individual colonies did not always receive the same number of workers that they produced. However, the number of workers introduced was adjusted to reflect the mean number of workers produced from focal colonies on each day of collection, allowing colonies to grow or shrink over time depending on the overall level of worker production. An exception to this was during Days 1-37 in Trial 1, when workers were instead introduced according to respective colony sizes to partially correct for the effects of mortality during this period. The introduced workers were randomised in relation to their colony of origin -- the proportion of workers from different colonies of origin was similar to the two treatments (Trial 1, *χ*^*2*^ = 19.9, *d.f.* = 14, *p =* 0.13; Trial 2, *χ*^*2*^ = 34.1, *d.f.* = 30, *p =* 0.28). Introduced workers were individually marked by unique colored numbered tags, or by non-unique tags/paint marks. All newly emerged workers could easily be identified by their reduced yellow pigmentation and the absence of tags/marks. The redistributions of workers continued throughout the experiment, until Day 54 (Trial 1) or Day 61 (Trial 2) of the experiment.

### Measuring Colony Performance

The following measurements were taken to assess colony performance (Figure 1). First, we recorded the total number, and the average size of newly emerging workers and males emerging in each colony. From this data we calculated the total number and average size of adults produced per colony (i.e. workers plus males). Second, the number of workers per colony was periodically assessed in a colony census. Given that we controlled the number of workers introduced into the colony, we used the census as a proxy of mortality. Censuses were conducted during the evening, when all or most foragers were expected to have returned to the nest. Third, the number of full nectar (syrup)-containing cells and pollen-containing cells was estimated every 1-3 days by visual inspection in the evening. These provide a measure of food collected and stored by each colony. Fourth, the data from foraging observations (see Behavioral Observations) allowed an estimate of each colony’s foraging rate, by calculating the rate (observations per minute) of individuals leaving or returning to each nest across all observation periods. In addition to the uniquely tagged workers, the foraging observations also included non-uniquely marked workers or, rarely, unmarked workers. Fifth, the accumulated productivity of the colony, as far as maintained until the end of the experiment, was assessed by weighing each colony’s nest comb (containing wax, silk cocoons, stored food and developing brood produced by the colony).

### Behavioral Observations

Starting on Day 34 (Trial 1) or Day 29 (Trial 2) and lasting until the end of the experiment, eight colonies (four per treatment) in each trial were regularly observed in 60 or 80 minute sessions, which quantified in-nest and foraging behavior of tagged workers in both treatments. In total, 40 in-nest sessions and 40 foraging sessions (6,400 min = 106.6 h total) were conducted in Trial 1, and 52 in-nest sessions and 52 foraging sessions (7,760 min = 129.3 h total) were conducted in Trial 2. This amounts to 236 hours of observations in total, across both trials.

In-nest observation sessions consisted of four 4-minute scans per colony, while foraging observation sessions consisted of two 20-minute scans per pair of colonies. During each in-nest observation scan, each visible tagged worker was watched once for 5-20 seconds, and its behavior was classified as one of the following: \**tending brood* (nursing); \**constructing*; *grooming*; \**fanning, feeding*, \**depositing food, egg-inspecting, aggression, walking, standing*, and, in Trial 2 only, \**incubating brood*. For detailed descriptions of each behavior, see Supplementary Methods. Of these behaviors, only those indicated with asterisks ‘*’ were considered ‘tasks’ that were later used to calculate the division of labor (see Statistical analysis and hypothesis testing).

During foraging scans, the behaviors observed were: \**leaving* = worker flying out from the end of the tube; *orientation =* leaving for orientation flight, distinguished by stereotyped slow circling flight when leaving the nest; \**returning with pollen =* worker with pollen in their pollen baskets landing on the platform and entering the nest via the tube (assumed to be returning from a pollen foraging trip); \**returning without pollen =* as above, but with no pollen seen (assumed to be returning from a nectar foraging trip). Events in which workers entered the tube, but flew back out without entering the colony, were not counted.

Behavioral observation data were used to compare the behavior of workers in each of the two treatments. In order to do this, the data were first cleaned in order to correct any likely tag identification errors made during recording (for details, see Supplementary Methods). The cleaned data from both types of scans were then used to create a behavioral profile for each worker, consisting of the frequency at which she was recorded performing each behavior over the course of the trial. Workers with fewer than five records in total (including “walking” and “standing”) were discarded for having insufficient data, giving final sample sizes of *n =* 251 in Trial 1, and *n =* 555 in Trial 2. The remaining workers that were used for our analyses had a mean of 19 (Trial 1) or 16 (Trial 2) records per individual. These behavioral profiles were then used for several types of analysis (Figure 1). Firstly, a foraging score for each worker was calculated by summing the frequency of ‘leaving’, ‘returning without pollen’ and ‘returning with pollen’. A nursing score was calculated by dividing the number of brood tending observations for a worker by its total number of in-nest records; this estimates the proportion of a worker’s in-nest time which was spent tending brood. Based on these measures, the ‘role’ of each worker was classified using arbitrary thresholds, as follows: ‘forager’ (foraging score > 4 AND nursing score < 0.5; Trial 1, *n =* 47; Trial 2, *n =* 84), ‘nurse’ (foraging score < 2 AND nursing score > 0.4; Trial 1, *n =* 71; Trial 2, *n =* 169) or ‘intermediate’ (all other workers; Trial 1, *n =* 133; Trial 2, *n =* 302).

### Statistical *Analyses and Hypothesis Testing*

Firstly, we evaluated the combined effect of treatment and worker behavior on colony performance across both trials, by modelling the effect of several factors on four different performance measures of each colony. Each of the four measures — estimated adult mass, mean number of nectar cells, mean number of pollen cells and comb mass at the end of each experiment — was used as the response variable in a separate model. Although several differences existed between trials, we partially account for this by using a linear mixed model with trial as random intercepts. The fixed predictors were: division of labor (DoL) metric, treatment, treatment x DoL metric, mean worker inactivity, standard deviation of worker foraging score, colony foraging rate and treatment x colony foraging rate. The DoL metric quantified the ‘division of individuals into tasks’ (Gorelick and Bertram, 2007; Gorelick et al., 2004), which approaches one when different workers specialize on different tasks, and is robust to differences in the number of workers between colonies (Gorelick et al. 2004). This was calculated based on all recorded ‘tasks’ (i.e. not all behaviors; see description of behaviors above) performed by individually-tagged workers. Colony foraging rate was calculated by dividing the total number of foraging observations for each colony by the total number of minutes spent observing that colony. This last measurement, unlike the DoL metric, mean inactivity and SD of foraging score, also included records for workers that were not individually tagged.

To investigate each trial (which differed in colony genotypes and the degree of genetic diversity, year and environmental conditions, supplemental food etc; Table S1) more specifically, the remainder of the analyses were performed separately for the two trials. We focussed on each hypothesis in turn.

Colony performance measures were compared between the two treatments in order to test the first hypothesis, that diverse colonies would outperform homogeneous colonies. We used Wilcoxon rank sum tests with continuity correction to compare nest comb mass and the total mass of adults produced over the whole experiment by colonies in the two treatments. Non-parametric tests were chosen for this and other per colony measures as a conservative analysis, because small sample sizes (e.g., number of colonies) make it difficult to confirm normality of data. To estimate total adult mass, we multiplied the number of adults by the cubed marginal cell length of each adult. We compared the difference in the number of workers over time between colonies in the two treatments using a linear mixed model (hereafter LMM), with response variable: number of workers (colony size) on census date; fixed predictor variables: treatment, day, and treatment x day interaction; and random intercept predictor variable: colony ID. We compared the level of stored nectar and pollen per colony over time between the two treatments using generalised linear mixed models (hereafter GLMMs) with a Poisson error distribution and with response variable: number of pollen cells or number of nectar cells; fixed predictor variables: treatment, day, and treatment x day interaction; and random intercept predictor variable: colony ID. We compared colony foraging rates between treatments using separate Wilcoxon rank sum tests for the total number of records of workers which were leaving or returning to the nest.

We then tested the second hypothesis, that workers in homogeneous colonies attempt to compensate for a lack of predisposed specialists. Firstly, to test Hypothesis 2a, which states that homogeneous colonies increase the production of small and large “predisposed specialists”, we compared the mean and standard deviation of worker size produced by each colony between treatments. We tested the effect of treatment within each trial using Wilcoxon Rank Sum tests, and the effect of treatment across both trials using linear mixed models, with random intercepts for trial. Secondly, to test Hypothesis 2b, i.e. whether middle-sized workers increase their level of specialization via behavioral plasticity, we first performed Spearman rank correlation to check the overall relationship between foraging and nursing, when pooling workers from both treatments together, and also to check the relationship between size and foraging: nursing ratio in each treatment. The level of division of labor between colonies in each treatment was compared with Wilcoxon Rank Sum test, using the DoL metric (explained above), calculated using either medium-sized workers or all workers. We next compared the proportion of ‘foragers’, ‘nurses’ and ‘intermediates’ among middle-sized workers between treatments, using Chi-squared tests. We also tested within medium workers in homogeneous colonies to check if they held roles in accordance with size, using one-tailed Wilcoxon rank sum tests to determine whether nurses were smaller than foragers. To test mean foraging and foraging specialization in workers more specifically, we compared the mean and variance of per-worker foraging score between treatments using Wilcoxon rank sum tests and Levene’s tests for homogeneity of variance, respectively. The same was measured for nursing score. Thirdly, to test Hypothesis 2c, i.e. whether workers altered their levels of inactivity, we compared the mean per-worker counts of inactivity between treatments using Wilcoxon Rank Sum tests, using either workers of all sizes, or only medium-sized workers.

Data processing and statistical analyses were performed using R (R Core Team 2017). In all linear models, the presented significance of the slope of fixed covariates (e.g. ‘day’) refers to the slope for the diverse treatment. The significance of a covariate x treatment interaction refers to the slope of the homogeneous treatment, as compared to the diverse treatment. For further details of analyses, see Supplementary Methods.

## RESULTS

In pooled analyses that included data from both trials and tested the effects of various factors on colony performance using mixed models, we identified several factors with a statistically significant effect on any of the four colony performance measures tested (Figure 2). Inactivity significantly decreased the total number of nectar cells (LMM, *n* = 19, *t* = -2.8, *p* = 0.03) and showed a statistically non-significant trend toward lower adult mass, comb mass and number of pollen cells. We also confirmed that any effect of inactivity was not mostly explained by the activity of specialists alone (see Supplementary Results and Table S2). Foraging rate significantly increased the estimated total adult mass (LMM, *n* = 19, *t* = 3.2, *p* = 0.014) and comb mass (LMM, *n* = 19, *t* = 4.44, *p* = 0.0004). Treatment (variation in body size distribution) also had a significant effect on comb mass, with diverse colonies being heavier (LMM, *n =* 19, *t* = -3.2, *p =* 0.006) and a non-significant trend for the number of nectar cells. Furthermore, the interaction between treatment and foraging rate showed a strong trend for an effect on comb mass, that almost reached our threshold for statistical significance, with foraging rate increasing the final comb mass with a steeper slope in homogeneous colonies (LMM, *n* = 19, = 3.8, *p =* 0.052). Thus, homogeneous colonies actually tend to produce smaller comb only when their foraging rate is low. None of these factors had a significant effect on more than one performance measure, and there were no significant effects of the other factors tested: DoL metric, standard deviation of foraging scores, or DoL metric x treatment interaction. In the following sections we compared the effect of treatment in each trial separately.

**Figure 2.**
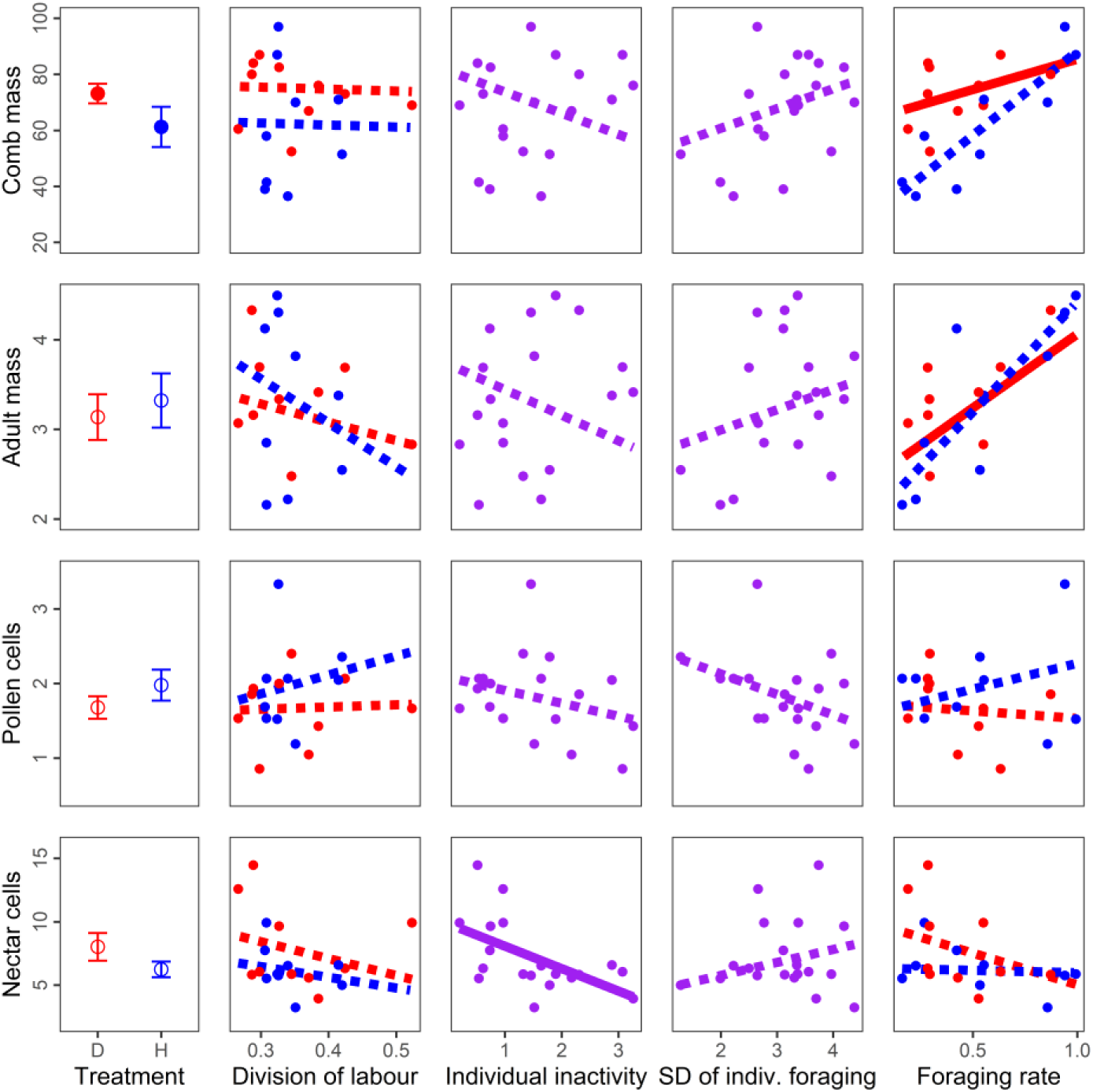
Plot matrix summary of linear models testing the effects of body size distribution and behavioral measures on indices of colony performance. Linear mixed models used a random intercept effect for trial (not shown), each with a colony performance index as a response variable (matrix rows), and a behavior or manipulation as a fixed predictor variable (matrix columns). Response variables were: nest comb mass at the end of the experiment (g), total adult mass (arbitrary units calculated by cubing the mean marginal cell length and multiplying it by the total number of adults produced for each colony), mean number of full pollen cells per census, mean number of full nectar cells per census. Predictor terms were: treatment, division of labour metric, division of labour x treatment interaction, mean inactivity per worker, SD of individual-level foraging, foraging rate, total foraging x treatment interaction. In terms where no interaction was tested, purple points show data and purple lines show linear model best fits. For interactions with treatment, red/blue points and lines show separate data and fits for the diverse/homogenous colonies respectively. Solid purple or red lines indicate significant terms in the linear mixed models (a solid blue line would indicate a significantly different effect for the homogeneous treatment). The leftmost column shows mean values for each treatment (red=diverse colonies, blue=homogeneous colonies), with solid circles indicating a significant effect of treatment.

### Colony Performance

In both trials, the estimated mass of adults produced per colony did not differ between treatments (Wilcoxon rank sum tests, Trial 1, *W =* 7, *n =* 9, *p =* 0.56; Trial 2, *W =* 21, *n =* 11, *p =* 0.329; upper row in Figure 3). Colony comb mass at the end of the experiment was similar between treatments in Trial 1 (Wilcoxon rank sum test, *W =* 11.5, *n =* 9, *p =* 0.81), but was significantly higher in diverse colonies in Trial 2 (*W =* 1, *n =* 11, *p =* 0.008, lower row in Figure 3).

**Figure 3.**
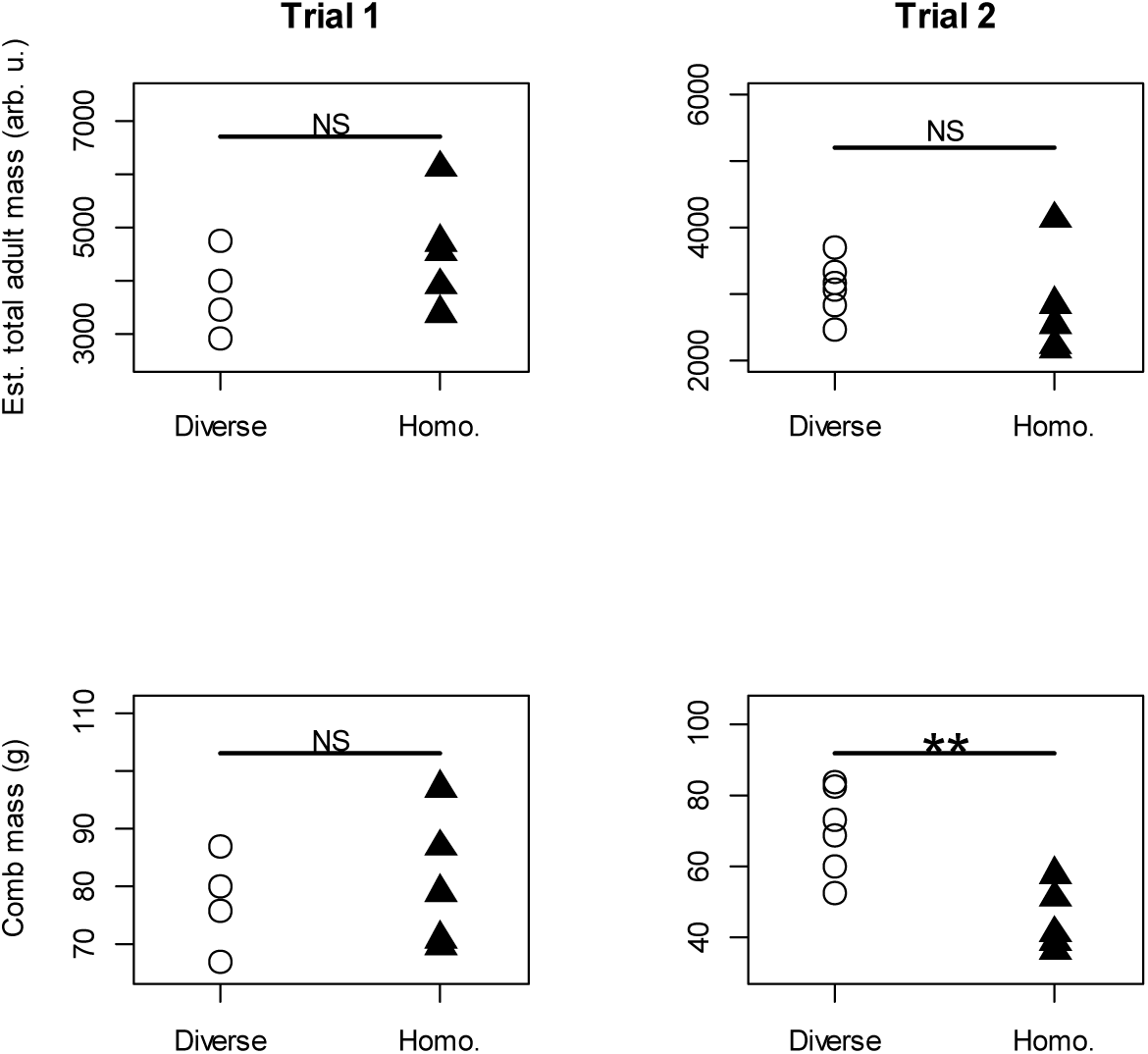
Colony performance measures as a function of worker body size distribution. Results shown for diverse or homogeneous (“Homo.”) colonies, separately for Trial 1 (left column) and Trial 2 (right column). Upper row: the total mass of adults produced over the course of the experiment per colony. Arbitrary units calculated by cubing the mean marginal cell length and multiplying it by the total number of adults produced for each colony. Lower row: nest comb mass per colony at the end of the experiment. Comb mass includes wax, silk cocoons, stored food and developing brood produced by the colony, but not adult bees. ** p < 0.01; NS = p > 0.05.

In Trial 1, worker number at the final part of the experiment was significantly affected by day, with worker number in diverse colonies getting smaller as the experiment progressed (LMM, *F* = 62.3, n = 54 colonies, *p* < 1×10^−6^; Figure 4), and also by a treatment x day interaction, with a greater decline over time in homogeneous colonies (LMM, *F* = 23.5, *p* < 1×10^−3^). In Trial 2, worker number was significantly affected by treatment, with diverse colonies being larger (LMM, *F* = 4.98, *n* = 110, *p =* 0.047; Figure 4) and by day, with worker number getting smaller as the experiment progressed (LMM, *F* = 72.3, *p* < 1×10^−12^), but not by a treatment x day interaction (LMM, *F* = 0.066, *p =* 0.80).

**Figure 4.**
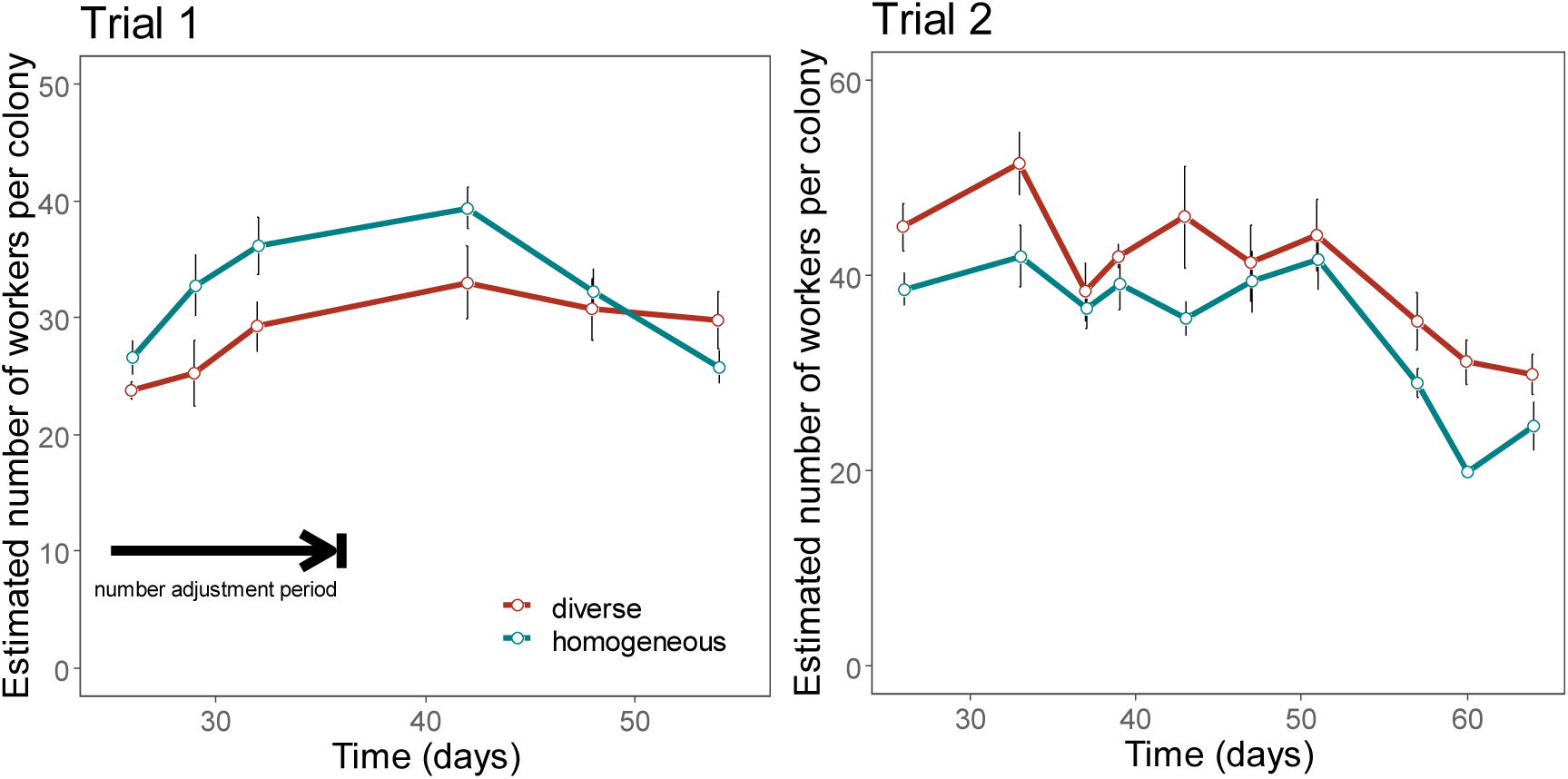
Estimated number of workers per colony over time. Trial 1 (Top) and Trial 2 (bottom). The plots show mean (±SE). Thick black arrow indicates a period in Trial 1 when workers were introduced depending on mortality levels in an attempt to keep colony size similar between treatments at this stage (there was no corresponding period in Trial 2). Other than this period, workers were introduced in equal numbers and so differences in colony size reflect differences in worker mortality. See Methods for additional details on census procedure and statistics.

The influence of body size distribution on colony food storage also differed between trials. In Trial 1, there was no effect of day on nectar cells in diverse colonies (poisson GLMM, *z* = 0.94, *p =* 0.35), but nectar cells in homogeneous colonies significantly declined more over time (treatment x day interaction, *z* = -4.00, *p* < 1×10^−4^; Figure S1). A post-hoc Wilcoxon rank sum test showed no significant difference in nectar cells between treatments on the final day of measurement (*n* = 9, *W* = 12, *p* = 0.52). In Trial 2 there was a significant increase by day in diverse colonies (poisson GLMM, *F* = 17.4, *p* < 1×10^−4^) and a significant interaction between day and treatment, with nectar cells increasing relatively less over time in homogeneous colonies (*F* = 22.3, *p* < 1×10^−5^). A post-hoc Wilcoxon rank sum test showed no significant difference in nectar cells between treatments on the final day of measurement (*n* = 11, *W* = 22, *p* = 0.23). In Trial 1, the number of pollen cells was not significantly affected by day in diverse colonies (zero-inflated poisson GLMM, *F* = 2.1, *p =* 0.15; Figure S1), but there was a significant day x treatment interaction, with pollen cells relatively increasing over time in homogeneous colonies (*F* = 6.4, *p =* 0.012). A post-hoc Wilcoxon rank sum test showed no significant difference in pollen cells between treatments on the final day of measurement (*n* = 9, *W*= 5, *p* = 0.35). In Trial 2, the number of pollen cells was not significantly affected by treatment (zero-inflated poisson GLMM, *F* = 0.01, *n* = 165, *p =* 0.93; Figure S1), by day (*F* = 1.5, *p =* 0.22), nor by a day x treatment interaction (*F* = 0.00, *p =* 0.97).

Colony foraging rates did not differ significantly between diverse and homogeneous colonies for either leaving the colony entrance (Wilcoxon rank sum tests; Trial 1, *W =* 3, *n =* 8, *p =* 0.2; Trial 2, *W =* 20.5, *n =* 11, *p =* 0.36), or returning to the colony entrance (Wilcoxon rank sum tests; Trial 1, *W =* 3, *n =* 8, *p =* 0.2; Trial 2, *W =* 18, *n =* 11, *p =* 0.66, Figure S2). For further details, see Supplementary Results.

### Responses to Reduced Size Distribution

We first assessed whether colonies with reduced worker size distribution compensate by producing larger or more size-diverse workers (*Hypothesis 2a*). There was no effect of treatment on the mean body size of workers emerging in each colony when testing each trial separately (Trial 1, Wilcoxon rank sum test, *W =* 9, *n =* 9, *p =* 0.55; Trial 2, *W =* 14, *n =* 11, *p =* 0.93, Figure S3 left column), or in a pooled analysis across both trials (LMM, *t* = *0.74, n* = 20, *p =* 0.47). There was also no effect of treatment on the body size standard deviation of workers emerging in each colony when testing each trial separately (Trial 1, Wilcoxon rank sum test, *W =* 20, *n =* 9, *p =* 0.15; Trial 2, *W =* 16, *n =* 11, *p =* 0.93; Figure S3 right column), or in a pooled analysis across both trials (LMM, t = 1.0, *n* = 20, *p =* 0.34).

We next tested whether size homogeneous colonies increase the proportion of specialists via behavioral plasticity (*Hypothesis 2b*). We first confirmed the presence of a typical division of labor with a negative correlation between the frequency of nursing and foraging observations per worker (Pearson’s correlation pooling both treatments; Trial 1, r = -0.25, *n =* 251, *p =* 6 x 10^−5^; Trial 2, r = -0.10, *n =* 555, *p =* 0.013; separately per treatment, see Supplementary Results, Figure 5 upper row), and a positive correlation between body size and the foraging: nursing ratio, in the diverse treatment (Spearman rank correlation; Trial 1, rho = 0.46, *n =* 119, *p =* 1 x 10^−7^; Trial 2, rho = 0.41, *n =* 217, *p =* 2 x 10^−12^; Figure 5 lower row). We found that the overall degree of specialization, as captured by the DoL metric, was similar in the homogeneous and diverse colonies in both trials, when restricted to comparing medium-sized workers (Wilcoxon Rank Sum tests, Trial 1, *n =* 8, *W =* 6, *p =* 0.69; Trial 2, *n =* 11, *W =* 15, *p =* 1), as well as when including all workers (Wilcoxon Rank Sum tests, Trial 1, *n =* 8, *W =* 6, *p =* 0.69; Trial 2, *n =* 11, *W =* 17, *p =* 0.79). Consistent with this analysis, in both trials, the proportion of medium sized workers which were ‘specialists’ (i.e. classified as nurses or foragers) was not significantly different between treatments (Chi-squared tests, Trial 1, *χ*^*2*^ = 4.0, *d.f.* = 2, *p =* 0.13; Trial 2, *χ*^*2*^ = 0.60, *d.f.* = 2, *p =* 0.74). Remarkably however, in both trials, foragers were significantly larger than nurses in homogeneous colonies (one-sided Wilcoxon rank sum tests; Trial 1, *W =* 577, *n =* 61, *p =* 0.027; Trial 2, *W =* 2388, *n =* 127, *p* < 0.001; Figure 6). When further testing for changes in foraging among medium sized workers, the mean per-worker foraging score did not differ between treatments (Wilcoxon rank sum tests, Trial 1, *W =* 2229, *n* = 168, *p =* 0.16; Trial 2, *W =* 16049, *n =* 385, *p =* 0.25; Figure 7 upper row). However, the variance of foraging score was greater in homogeneous colonies in Trial 1 (Levene’s test, F = 13.2, *n* = 168, p < 0.001), but not in Trial 2, (Levene’s test, F = 0.71, *n =* 385, *p =* 0.40). The effect of treatment on per-worker nursing score also differed between trials (see Supplementary Results and Figure S4).

**Figure 5.**
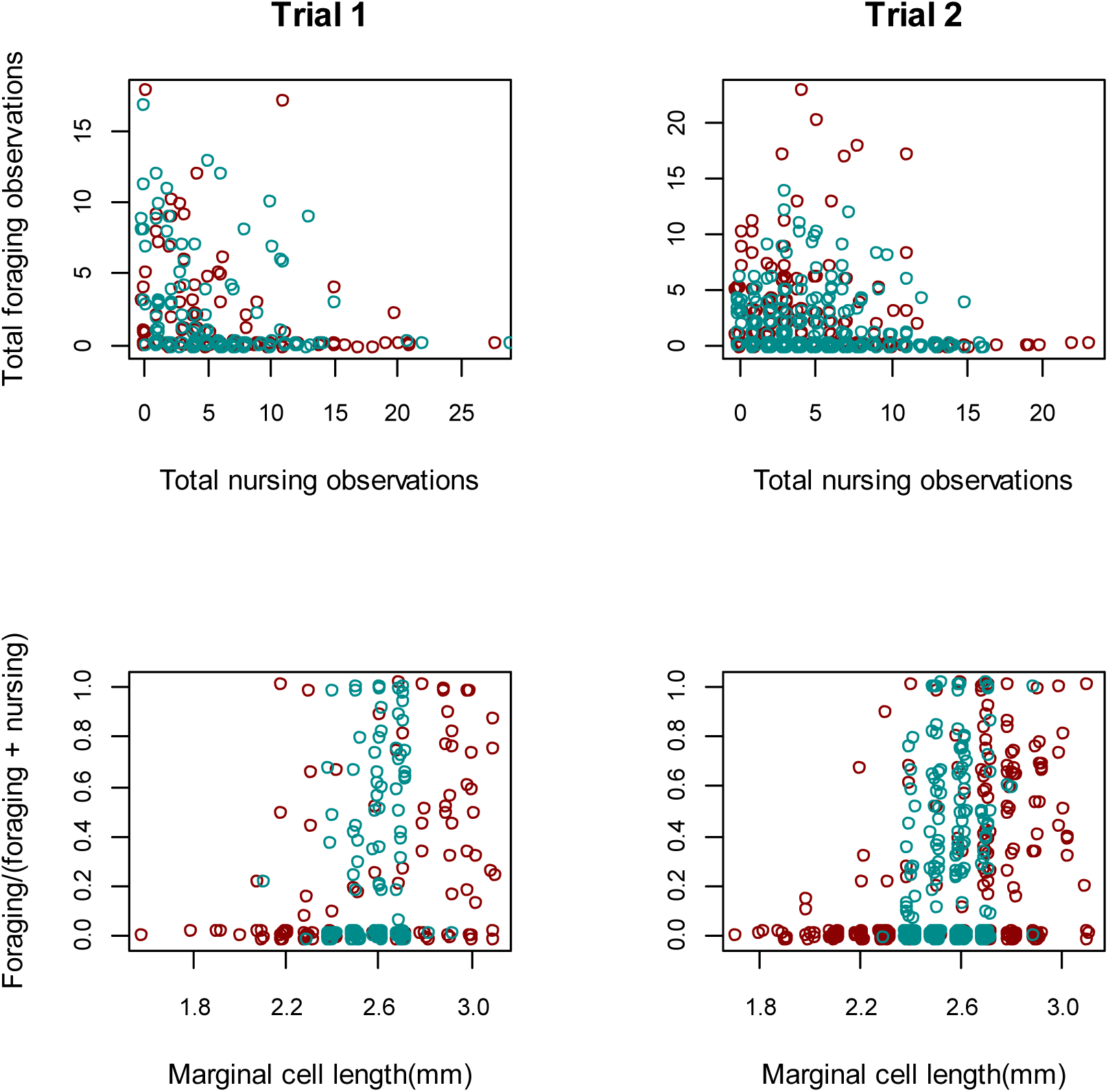
Nursing and foraging performance in size–diverse and -homogenous colonies. Red symbols-individuals in diverse colonies; blue symbols – individuals in-homogeneous colonies. Trial 1 - left column; Trial 2 - right column. Upper row: significant negative correlations between total foraging observations and total nursing (brood tending) observations per worker. Lower row: foraging: nursing ratio as a function of worker body size (wing marginal cell length). In order to display multiple data points in the same position, a small amount of horizontal and vertical random noise has been added to each data point.

**Figure 6.**
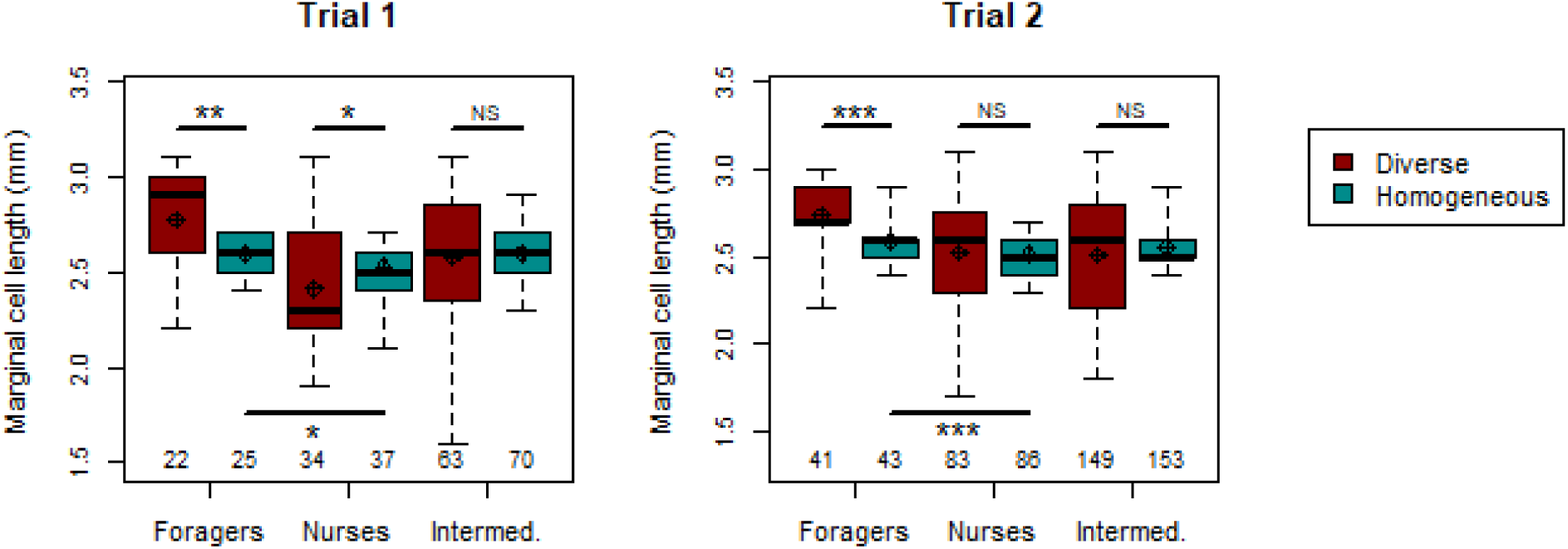
Body sizes of worker roles in size-diverse and -homogeneous colonies. Shown separately for Trial 1 (left) and Trial 2 (right). Workers categorised as foragers, nurses or intermediate, based on task frequencies (see Methods for details). For each trial, sample sizes of workers in each role and treatment shown at the bottom of the plot area. Diamonds = means; thick black lines = medians; boxes = interquartile ranges; dashed whiskers = ranges. Upper statistical comparisons based on individual Wilcoxon rank sum tests comparing the sizes of workers in each role between treatments. Lower statistical comparisons based on one-sided Wilcoxon rank sum tests comparing whether foragers were larger than nurses in the homogeneous treatment. *** p < 0.001; ** p < 0.01; * p < 0.05; NS = p > 0.05.

**Figure 7.**
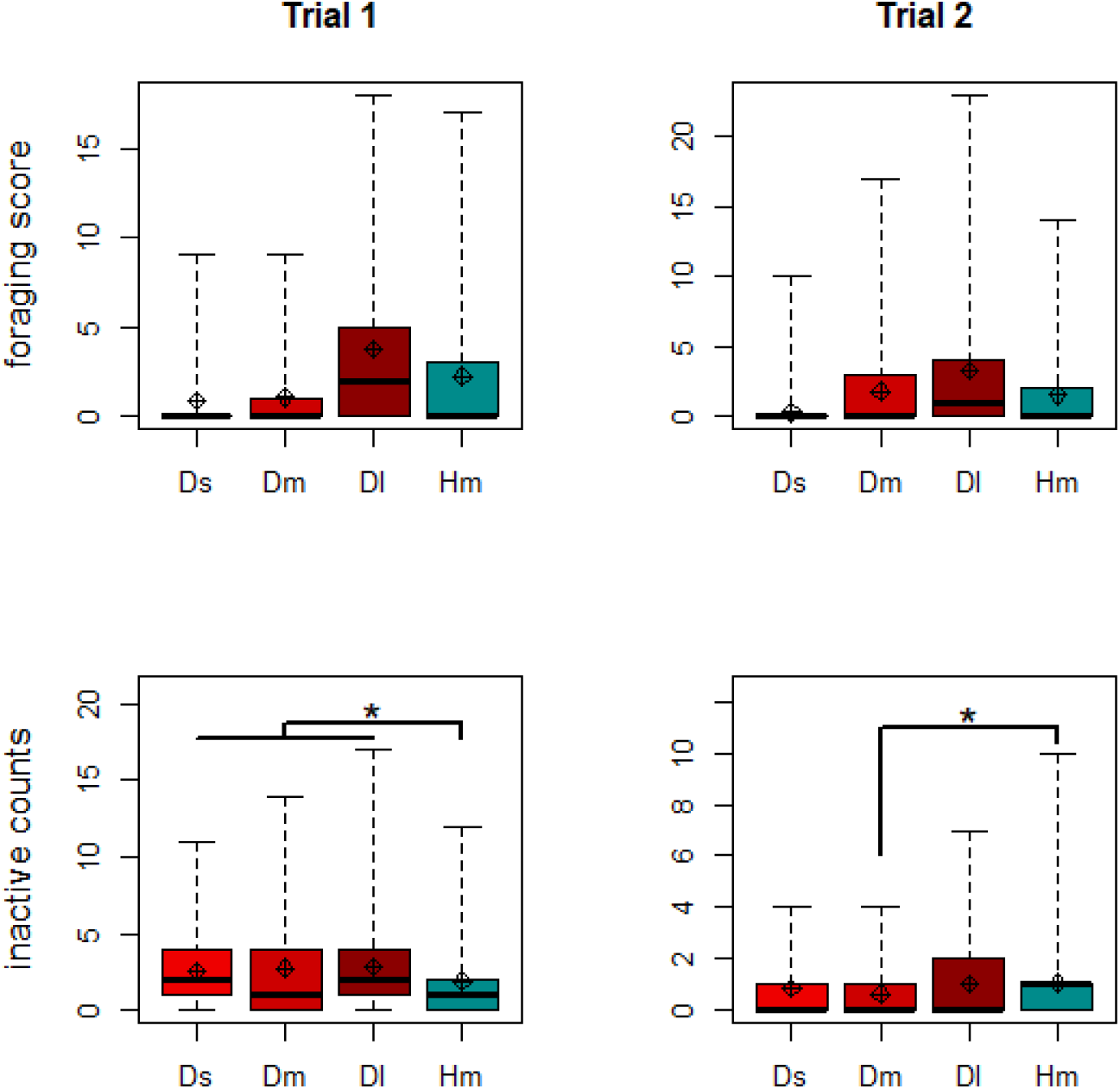
Worker foraging and inactivity as function of body size and colony size diversity. Shown for Trial 1 (left column) and Trial 2 (right column). Foraging score in upper row calculated by summing records for leaving the nest, returning to the nest without pollen and returning to the nest with pollen (see Figure S2 for more details). Inactivity counts in lower row based on records of workers standing with no obvious task. Ds = small workers from diverse colonies; Dm = medium workers from diverse colonies; Dl = large workers from diverse colonies; Hm = medium workers from homogeneous colonies. Diamonds = means; thick black lines = medians; boxes = interquartile ranges; dashed whiskers = ranges.

Another form of individual plasticity relates to the time spent inactive (Hypothesis 2c). In Trial 1, the mean per-worker level of inactivity (observations when a worker was seen standing still, with no obvious task) was significantly lower in homogeneous colonies (Wilcoxon rank sum test, *W =* 9028, *n* = 251, *p =* 0.037; Figure 7 lower row), but there was only a non-significant trend when restricting the comparison to only middle-sized workers (*W =* 2834, *n* = 168, *p =* 0.295). Given a lack of correlation between size and inactivity in the diverse colonies (Spearman rank correlation, rho = -0.02, *p =* 0.8), it is possible that the lack of a significant effect when restricting the comparison to middle-sized workers is due to reduced sample size and statistical power. In Trial 2, there was no significant effect of treatment when comparing the amount of inactivity for all workers in the colony (Wilcoxon rank sum test; *n =* 558, *W =* 37741, *p =* 0.802), but inactivity was higher in the homogeneous treatment when limited to only middle-sized bees (*n =* 385, *W =* 12508, *p =* 0.017; Figure 7).

## DISCUSSION

Previous studies suggest that specialization and/ or plasticity are strategies to improve collective behavior, including in social insect colonies. However, these strategies have been typically studied in different species (e.g., leaf cutter ants show profound morphological specialization, and honey bees profound behavioral and physiological plasticity) and their interaction has rarely been tested by using experimental manipulations under ecologically relevant conditions. We studied the interplay between these two strategies in bumble bee colonies freely-foraging under challenging environmental conditions. This ecological context forced them to collect resources and presumably contend with competitors, predators and parasites. Our findings provide the first evidence under field realistic conditions suggesting that both size-related specialization and individual plasticity are functionally significant in a bumble bee. Thus, the importance of body size variability is not limited to vast colonies of ants and termites. Plasticity in individual behavior is important in bumble bees and can at least partially compensate for the lack of large- or small-sized specialists under at least some conditions.

We first tested the commonly held hypothesis that predisposed specialization, as manifested in differently sized bumble bee workers, improves colony performance. This hypothesis predicts compromised performance in colonies that are restricted to middle-sized, presumably less specialized, workers. Taking the two trails together, colonies with typical body size diversity indeed did better in some indices of performance (Figure 2), lending credence to this hypothesis. However, a closer look at each trial separately reveals that size diverse colonies actually did better only in Trial 2, in which diverse colonies had significantly greater comb mass (Figure 3), more workers (Figure 4), and a relatively greater increase in number of nectar cells over time (Figure S1). These findings, which suggest an adaptive role for large and/ or small workers as predisposed specialists in bumble bees, are different from a laboratory study in which body size diversity did not affect performance in *B. impatiens* (Jandt and Dornhaus 2014), but are consistent with the ubiquity of body size -task association reported in bumble bees and also found in the current study (Figure 5). Remarkably, even within the reduced size range of the homogeneous colonies, the foragers were larger than the nurses (Figure 6). This overall effect of size diversity fits well with previous research showing that large workers have a range of traits which may make them better suited for foraging, such as larger eyes, larger brains and stronger circadian rhythms (see Introduction). The effect of reducing body size variability was however, smaller than we expected; homogeneous colonies in Trial 1 did no worse than diverse colonies in any measure, and may have even performed slightly better in some measures (e.g. Figures 4, 5 and S1). The cross-trial analyses further suggest that factors other than body size variability are also important for colony performance (Figure 2). These include decreased worker inactivity and increased foraging rate (particularly for homogeneous colonies). Thus, we found an effect of size-based specialization on colony performance, but its contribution was limited and context dependent. It is also worth noting that we had a relatively modest sample-size for between-colony comparisons and so our statistical power was limited in detecting nuanced effects.

The second hypothesis we tested was that compensation for reduced body-size diversity would occur by plastic responses at the colony or individual level. We found no evidence for influence of worker body size diversity on the sizes of new bees reared by the colony (*Hypothesis 2a*). Homogenous colonies did not produce larger workers or a broader worker body size range, in either trial (Figure S3). This finding for the bumble bee is different from some ant and termite species where morphologically distinct specialized soldiers are produced in response to colony needs (e.g. Chouvenc et al., 2015; Passera et al., 1996). Our findings are more consistent with plastic responses in the behavior of individual adult bees, involving changes in both specialization and effort (*Hypotheses 2b* and *2c*). In Trial 1, in which homogeneous colonies performed similarly to diverse colonies, medium-sized workers had greater variance in foraging than the same sized workers in diverse colonies (Figure 7), consistent with increased specialization or increased effort by specialists. Furthermore, in Trial 2, medium-sized workers in homogeneous colonies showed both a significantly higher level of nursing and a significantly higher variance in nursing, when compared to medium-sized workers in diverse colonies (Figure S4). Nevertheless, increased specialization cannot fully account for the improved performance of some homogeneous colonies. For example, medium-sized workers in homogeneous colonies were not more likely to be categorized as ‘nurses’ or ‘foragers’ compared to same size bees in typically diverse colonies, and the DoL metric was similar for diverse and homogeneous colonies, even when restricting the analysis to only medium sized bees. In addition, in the analyses across trials, the DoL metric was not significantly related to any measures of colony performance (Figure 2), with trends not even in the expected direction. Given that the contribution of increasing specialization to the performance of homogeneous colonies was limited, how were some of these colonies able to perform as well or even better than diverse colonies? We suggest that a significant factor contributing to their relatively good performance was an increase in the effort of individual bees.

In Trial 1, workers in homogeneous colonies spent significantly less time inactive compared to those in diverse colonies. The results analysed across both trials (Figure 2) further corroborated the importance of inactivity by showing a significantly negative effect of inactivity on the number of nectar cells and a statistically non-significant trend for the same effect on adult mass and number of pollen cells. Reducing the amount of inactivity has previously been suggested as a mechanism to buffer perturbations to colony composition (Charbonneau et al., 2017; Hasegawa et al., 2016). When we used the Aikake’s Information Criterion (AIC) to assess whether any effects of inactivity on performance was best explained by the activity of specialists (i.e., workers classified as nurses, foragers or both; Table S2), we found a nuanced effect at best. This suggests that increased effort by generalists may be equally or more important than increased effort by specialists or increased specialization. Nevertheless, a colony level increase in foraging effort contributes to the success of homogeneous colonies as suggested by the steeper (i.e. more positive) slope relating colony foraging rate to comb mass relative to diverse colonies (Figure 2, top right panel). The most successful homogenous colonies, all of which belonging to Trial 1, showed a very high rate of foraging. This analysis is consistent with medium-sized workers being less efficient foragers than large workers, and therefore needing to increase their foraging rate to a greater extent in order to achieve a similar comb mass. Higher foraging rate also increased the total adult mass produced, when analysed across trials (Figure 2). Taking these analyses together, we suggest that the decrease in inactivity and the increase in individual effort at least partially compensated for the lack of large and (perhaps) small specialists in homogeneous colonies, explaining the comparable performance of homogeneous and diverse colonies in Trial 1.

The slight increase in specialization and the increase in individual effort discussed above are similar to the short term increases in foraging or brood care efforts reported for workers in bumble bee colonies in which foragers (Crall et al. 2018), or nurses (Pendrel & Plowright 1981), were experimentally removed, respectively. Such an increase in individual effort is probably associated with an increase in energy consumption and in predation risk (for the foragers) and therefore may be costly. This high cost may explain why individual plasticity could not compensate for the lack of predisposed specialists in Trial 2. In addition, in Trial 1, the slightly higher number of workers in homogeneous colonies significantly declined towards the end of the experiment (significant interaction between treatment and time; Figure 4), with a similar trend apparent for the number of nectar cells (Figure S1). These findings may suggest that even in Trial 1, the ability to compensate for the lack of predisposed specialists was limited and declined towards the end of the experiment.

The reason why colonies responded differently between the two trials is not clear. As detailed in the Methods and Supplementary Methods, the two trials differ in several ways (summarised in Table S1). This includes the source colonies (that may differ genetically between the two trials), the degree of genetic variation among queens within trial, and the environmental conditions in the two years. Identifying the specific environmental or internal (including genetic) variables that contributed to the different results in the two trials is very challenging and beyond the scope of the current study. However, this variability suggests that multiple factors affect colony level performance, and that a similar experiment under different conditions may produce somewhat different results. For example, we assume that the effect of body size diversity may be more pronounced under conditions of lower ambient temperature and shorter days, under which large bees may be more efficient when foraging outside (Heinrich, 1979; Spaethe and Chittka, 2003). From a broader perspective, the advantage of size-based specialization under different environments has only been partially investigated, and is not yet fully understood (Couvillon et al., 2010; Peat et al., 2005; Kerr et al., 2019; Chole et al. 2019). We hope that our study will encourage additional studies under diverse ecologically relevant conditions.

A somewhat unexpected and remarkable finding of our study was that even within the limited size range of medium size workers in the homogeneous colonies, foragers were nevertheless significantly larger than nurses in both trials (Figure 6). This appears to show that when the smallest and largest workers are lost (or in our experiments, replaced), the new smallest and largest workers are more likely to take on the roles of nurses and foragers, respectively. This finding suggests that medium-sized workers can serve as a “buffer group” because they are best positioned to plastically adjust their level of specialization along with colony needs. This finding also raises an interesting question: how do workers adjust their behavior in accordance with relative body size? Although currently there is no clear answer to this question, several possibilities exist. Firstly, body size could affect an individual’s response threshold to task-related external stimuli, such that, for example, the larger the individual, the more likely it is to respond to foraging-related stimuli. In this case, a loss of large workers would result in levels of a stimulus rising until it reached the threshold of the new largest individuals, which would then begin foraging. Similar response threshold models for foraging have been suggested for honeybees (e.g. Fewell and Winston, 1992; Page et al., 1998; Sagili and Pankiw, 2007 and references therein) and ants (Robinson et al., 2009). A second possible explanation is that workers can assess their own size relative to that of their nestmates. Other organisms have been shown to assess the relative sizes of conspecifics by using tactile (Wells, 1988) or audible (Robertson, 1986) cues (visual cues can probably be excluded within the dark nests of bumble bees). The response threshold and assessment models could also operate in tandem, because an individual’s internal response threshold could itself be modulated in response to an assessment of its social environment. Similar mechanisms, such as learning-based changes in responsiveness, have been shown in other contexts in social insects, including bumble bees (reviewed in Jeanson and Weidenmuller, 2014). Worker body size is also related to other aspects of individual variation, e.g. spatial location within the nest, which has been shown to affect the propensity of workers to increase foraging activity after the removal of foragers from *Bombus impatiens* colonies (Crall et al., 2018). The mechanisms used by workers to plastically adjust their behavior in response to changes in colony composition deserves further investigation.

In conclusion, our study suggests that in the same species, both predisposed specialization and plasticity in the behavior of individuals can contribute to group performance in an ecologically-relevant context. We showed that a wide distribution of worker body sizes can improve colony-level performance in bumble bee colonies, which are relatively small in number. This finding is not consistent with the common view that size diversity is functionally significant only in large colonies with morphological castes. We also show that, even when size diversity is severely reduced, colonies do not always perform worse, even under extremely challenging environmental conditions as in our experiment. Our data are consistent with the premise that such compensation at the group level is at least partially achieved by flexibility in the behavior of individual workers. The results of our experiments begin to unravel the way that group systems can evolve to take advantage of both specialization and plasticity towards improving collective performance under the complex fluctuations of the natural environment. Understanding the interplay between these two mechanisms, is crucial for understanding how collective behavior is organized to appropriately respond to complex ecological conditions.

## Supporting information

Supplementary Methods & Results

## ACKNOWLEDGMENTS

We are grateful to the support of students and researchers in the research group of Guy Bloch, especially to the following, who assisted with behavioural observations in these experiments: Molly Avidan-King, Hadar Baron, Yogev Herz and Inbal Tziperman. We also wish to thank reviewers for their comments. This research was supported by grants from the National Academies Keck Futures Initiative (NAKFI, to ON, GB, and MP), the US-Israel Binational Science Foundation (BSF-NSF in Biology (ICOB) 2012807 to GB), the US–Israel Binational Agricultural Research and Development (BARD) fund (IS-4418-11 to GB), and a Lady Davis Fellowship held by JGH

