## Supplementary Methods & Results for "Is diversity in worker body size important for the performance of bumble bee colonies?"

*Colony foundation and male production*

In Trial 1, colonies were founded by unrelated queens that did not undergo diapause. These queens underwent two $CO_2$ narcosis treatments which circumvent diapause (Roseler, 1985; Tasei, 1994) and were provided with a conspecific 'stimulatory' worker in order to stimulate them to found colonies in the rearing facility. During this trial, several colonies produced males on or before day 25 of the experiment (n = 37 males across 7 colonies), meaning that they were produced from eggs laid before treatment began, given that the normal egg to adult developmental time is around 26 days (e.g. Bloch 1999; Duchateau and Velthuis 1988). Early male emergence is common in industry-produced colonies in which the queens do not experience natural diapause (Velthuis 2002). It is also possible that the introduced stimulatory workers laid eggs. We excluded the possibility that these males were caused by inbreeding, which can result in diploid males occurring (Duchateau et al. 1994), because the ratio of males to workers in the first 25 days was significantly different to 1:1, which is typical for colonies with diploid males (goodness of fit tests, all p-values $< 0.01$ across colonies; one colony produced ~80% males within this time period and the others produced 10% or less). Note that, as mentioned in the Methods (*Colony Maintenance and Treatment*), these early produced males were replaced with workers, which should have mitigated any detrimental effects of male production (i.e. males acting as a burden on colony resources). The colony which produced a high proportion of early males went on to produce mostly females later on.

In the second trial, we undertook several measures to reduce genetic variation among worker bees and to reduce early male production. Specifically, we used colonies that were all headed by sister-queens (i.e., emerging from a single mother colony) which were mated by brothermales (from another unrelated colony). This should mean that workers across different colonies had an average relatedness value of 0.625, which is almost as high as workers within the same colony (r = 0.75). We reasoned that this might help to control for variation between colonies in performance, aside from the effects of treatment. The queens also underwent a short (3-week) diapause (which more closely simulates natural conditions) before $CO_2$ treatment, and the conspecific stimulatory workers were removed or replaced (as necessary) after 1 week. These procedures substantially reduced the emergence of early males in this trial ($n$ = 1 male from a single colony).

In Trial 1, the mean eclosion date of the first worker (before receipt of colonies from supplier) was estimated to be four days before receipt of the colonies, i.e. 10th May 2015 (given that the number of workers upon receipt of the colonies was similar to Trial 2, where the mean first workers eclosion was known to have occurred four days before receipt). In Trial 2, the mean first worker eclosion date was known to be 22nd June 2016. Thus, four days should be added to the reported experimental days to calculate the date from first worker eclosion (in both trials).

The differences between trials are summarised in Table S1.

Table S1. Summary of differences between the trials.

| | **Trial 1** | **Trial 2** |
|---|---|---|
| **No. Colonies** | 9 | 11 |
| **Colonies closely related** | No | Yes |
| **Colony foundresses experienced diapause** | No | Yes |
| **Dates** | May-July 2015 | June-August 2016 |
| **Dead workers replaced periodically** | Only during the initial part of experiment | No |

*Behavioral classification*

Detailed descriptions of in-nest tasks are as follows: **tending brood* = sustained inspection of brood cells with mandibles and antennae, or feeding of larvae (type of brood tended, larva or pupa, was also recorded in Trial 2); **constructing* = the manipulation by mandibles of wax substrate, either by building/ modifying food cells, or by forming connections between food/brood cells, or otherwise the manipulation of other nest substrate (pulp from cardboard base of nest – recorded separately in Trial 2); *grooming* = sustained self-grooming of legs, thorax, head or proboscis; **fanning* = sustained (presumed thermoregulatory) fanning of wings, whilst standing still and raised on legs, usually standing on brood comb or wall of nestbox (and not directed towards other individuals); *feeding* = feeding self with proboscis in food cell; **depositing food* = obviously depositing nectar or pollen into food cell; *egg-tending* = sustained inspection of egg cells, eating eggs or laying eggs (recorded separately); *aggression* = agonistic behavior towards other workers or the queen (buzzing, antennating, darting, mandibulating/ biting or stinging attempts –each recorded separately); *walking* = actively moving, but with no obvious or sustained other behavior, often characterised by moving swiftly over the nest comb or around the periphery of nest, and only stopping briefly

(<2 seconds) to inspect food or brood cells; *standing* = stood still with no clear other behavior, often characterised with head in down position but with mandibles or antennae not moving, commonly consistent with sleep (but at other times apparently alert). In Trial 2 only, **incubating brood* was also distinguished, characterised by standing with body tightly pressed against brood and with abdomen pulsating.

*Data cleaning*

Occasionally, errors or uncertainties in tag observation records stemmed from difficulty seeing the exact tag ID, or from human error (e.g. recording the wrong color or number). This represented 8.8% and 5.1% of the total records in Trials 1 and 2, respectively. In these cases it was important to assign the tags to the most likely candidate tag ID where possible, using strict criteria (see below), in order to ensure the data were not biased against certain tags or certain behaviors. If no credible alternative tag identity was clear, or if two or more tags were equally credible, the tag record could not be reinterpreted and was discarded from the dataset. This was the case in 2.8% and 2.8% of the total records in Trials 1 and 2, respectively. Candidate tag credibility was determined by several parameters: the candidate tag ID must have been introduced into the focal colony, OR must have been an established drifter to the focal colony. In addition, the tag must have either been noted as a possible alternative at the time of recording, OR must have been a color-based mistake (i.e. with the same number), OR must have been similar in each of its digits (e.g. a 3 is mistakable for an 8). Any tag identities which were spotted in the same in-nest scan as the focal tag were assumed not to be a credible alternative (since a tag should only have been recorded once per in-nest scan). A tag identity which was recorded in other in-nest scans during the same session or day was taken as evidence in support of the tag being credible, because it showed that that bee was present at around the same time as the focal bee. The same is true for tag identities recorded during a single foraging scan (since multiple records of a forager were permitted during foraging scans, and foraging workers often foraged in back-to-back foraging bouts).

*Statistical modelling*

For LMMs and GLMMs, the 'lmer' or 'glmer' functions from R package 'lme4' (Bates et al. 2015), or the 'glmmadmb' function from the R package 'glmmADMB' (Fournier et al. 2012; Skaug et al. 2013) were used. GLMMs were fitted by maximum likelihood using Laplace approximation. In all linear models, non-significant fixed effects were removed from the model in reverse significance order (interaction terms before subsumed main effects), with p-values determined by likelihood ratio tests. For any fixed effects remaining in the minimally adequate model (i.e. where all terms significant), reported p-values are the result of Wald z tests. Following guidance from Bolker et al. (2009), Wald F or Wald t tests were used instead where overdispersion was present, via the lmer function or the Anova function in the R package 'car'. In all models, the significance of fixed factors refers to the difference in intercept between the two factor levels (e.g. homogeneous vs. diverse treatment). The significance of the slope of fixed covariates (e.g. 'day') refers to the slope for the diverse treatment. The significance of a covariate x treatment interaction refers to the slope of the homogeneous treatment, as compared to the diverse treatment. For example, the result of a non-significant effect of day, but a significant day x treatment interaction, means that the slope of day for the diverse treatment was not significantly different from zero, but the slope of day for the homogeneous treatment was significantly different from the slope of day for the diverse treatment.

In the models for comparing the number of food cells, generalised linear mixed models (GLMMs) were used with a Poisson error distribution, which is appropriate for count data as a response variable. A log link function was used, since this was found to produce a more normal error distribution, when compared to an identity link function (i.e. without transformation) in each model. Day was included as the number of days since ad libitum feeding ceased (i.e. the intercepts were fixed at this time), since modelling the effect of day on food cells before this time was considered nonsensical for interpretation. For the pollen models, zero-inflated GLMMs were used (with the glmmADMB package) to account for the large number of zeros in the pollen cell count data. Where there was a significant treatment x day interaction, the effect of treatment on the final day of measurement was assessed using a post-hoc Wilcoxon rank sum test.

### SUPPLEMENTARY RESULTS

*The inactivity of specialists*

We checked whether any effects of (in)activity on performance were mostly mediated by the activity of specialists. In each of the four performance models (combining the data from both trials), we tried substituting our original inactivity term (the mean inactivity of all workers) with three alternative inactivity terms: 1) mean inactivity of foragers, 2) mean inactivity of nurses, and 3) mean inactivity of specialists (foragers + nurses). To do this, we used simple versions of the models with inactivity as the only fixed predictor. We were then able to compare the Aikake's Information Criterion (AIC) scores for the models using each term, which revealed whether using these terms provided a better explanatory power (lower score). However, in all but one (very marginal) case, the AIC values of models with the alternative terms were not lower than models with the original term (see analysis table below). Thus, it seems the mean activity of specialists was not a better predictor of colony performance than the mean activity of all workers, i.e. not only the activity of specialists was important

Table S2. AIC values comparing different inactivity terms for simplified versions of each performance model, i.e. lmer (metric ~ inactivity + (1|trial)). Bold values indicate the model with the lowest AIC for each performance metric).

| Inactivity term used | Adult Mass | Comb Mass | Nectar Cells | Pollen Cells |
|---|---|---|---|---|
| **Mean inactivity of all workers** | **289.8** | 153.6 | **92.5** | **41.5** |
| **Mean inactivity of foragers** | 290.2 | 154.5 | 95.8 | 42.9 |

| | | | | |
|---|---|---|---|---|
| **Mean inactivity of nurses** | 290.5 | 155.1 | 95.1 | 42.6 |
| **Mean inactivity of foragers and nurses** | 289.9 | **153.5** | 94.8 | 42.4 |

*Pollen and nectar foraging rates*

Further to comparing the rate of leaving the nest to forage and returning to the nest (see Results), we also compared the rate of bees returning to the nest with or without (assumed to be mostly nectar foraging) pollen. For both measures, there was no statistically significant difference between treatments (Figure S2; Wilcoxon rank sum tests; without pollen: Trial 1, $W = 2.5$, $n = 8$, $p = 0.15$; Trial 2, $W = 16$, $n = 11$, $p = 0.93$,; with pollen: Trial 1, $W = 3$, $n = 8$, $p = 0.2$; Trial 2, $W = 22$, $n = 11$, $p = 0.23$; Figure S2).

*The correlation between nursing and foraging activity of individual bees*

When analysing each treatment separately, a significant negative correlation between the frequency of nursing and foraging observations per worker was found for the two treatments in Trial 1 (diverse treatment, r = -0.25, $n = 119$, $p = 0.006$; homogeneous treatment, r = -0.25, $n = 132$, $p = 0.004$), while for Trial 2 the relationship fell short of significance (diverse treatment, r = -0.11, $n = 273$, $p = 0.068$, homogeneous treatment, r = -0.10, $n = 282$, $p = 0.103$).

*The body sizes of nurses and foragers across treatments*

In both trials, workers classed as ‘foragers’ were significantly larger in the diverse treatment than in the homogeneous treatment (Wilcoxon rank sum tests; Trial 1, W = 421, n = 47, $p = 0.002$; Trial 2, W = 1740, n = 92, $p < 1 \times 10^{-8}$; Figure 7). Workers classed as ‘nurses’ were smaller in the diverse treatment than in the homogeneous treatment in Trial 1 (Wilcoxon rank sum test; W = 437, n = 71, $p = 0.026$), but not in Trial 2 (Wilcoxon rank sum test; W = 3710, n

= 167, *p* = 0.46). In both trials, body size of bees classed as 'intermediate' was similar for bees in the diverse and homogenous treatment colonies (Wilcoxon rank sum tests; Trial 1, W = 2121, n = 133, *p* = 0.70; Trial 2, W = 10929, n = 299, *p* = 0.74).

*Nursing levels*

The nursing count (tending larvae or pupae) of medium-size workers Trial 1, was not significantly different between treatments (Wilcoxon rank sum test, W = 2545, *p* = 0.86), but the variance was slightly but significantly smaller in homogeneous colonies (Levene's test, F = 5.4, *p* = 0.02;  Figure S4a). In Trial 2, both the per-worker nursing count (Wilcoxon rank sum test; n = 387, W = 12750, *p* = 0.012; Figure S4b), and the variance (Levene's test; n = 387, F = 4.56, p = 0.033) were significantly higher in the homogeneous treatment.

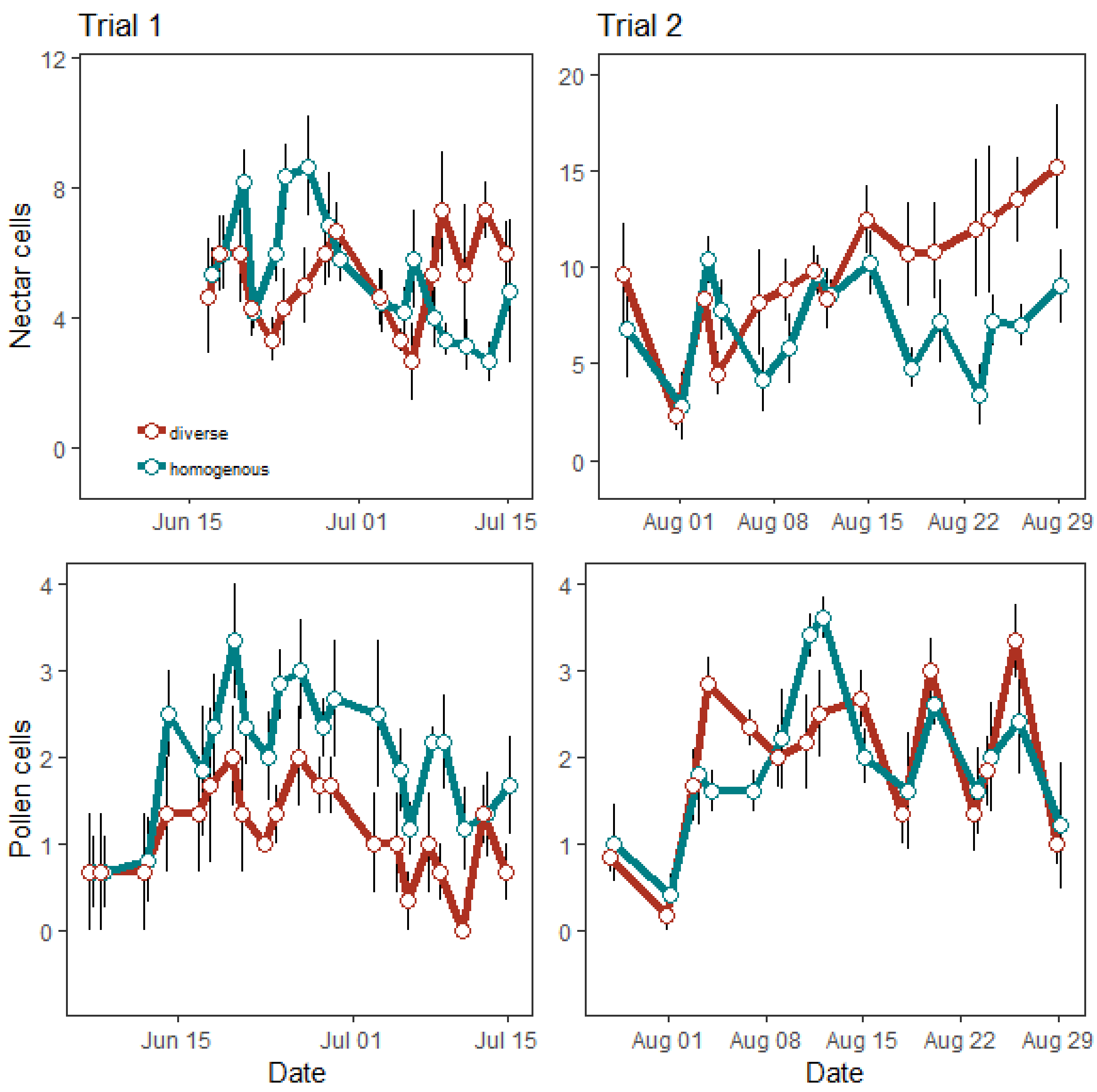


**Figure S1**. Estimated number of full pollen and nectar cells over time in size-diverse and -homogeneous colonies. Shown are mean ± SE for Trial 1 (left column) and Trial 2 (right column). The numbers were estimated based on visual inspection of cells each evening. Colonies were connected to the outside environment on 25 or 26 May 2015 (Trial 1) or on 12 July 2016 (Trial 2). In top left plot, the blank area before 17 June indicates period when some supplemental syrup remained in colonies and could not easily be distinguished from collected nectar.

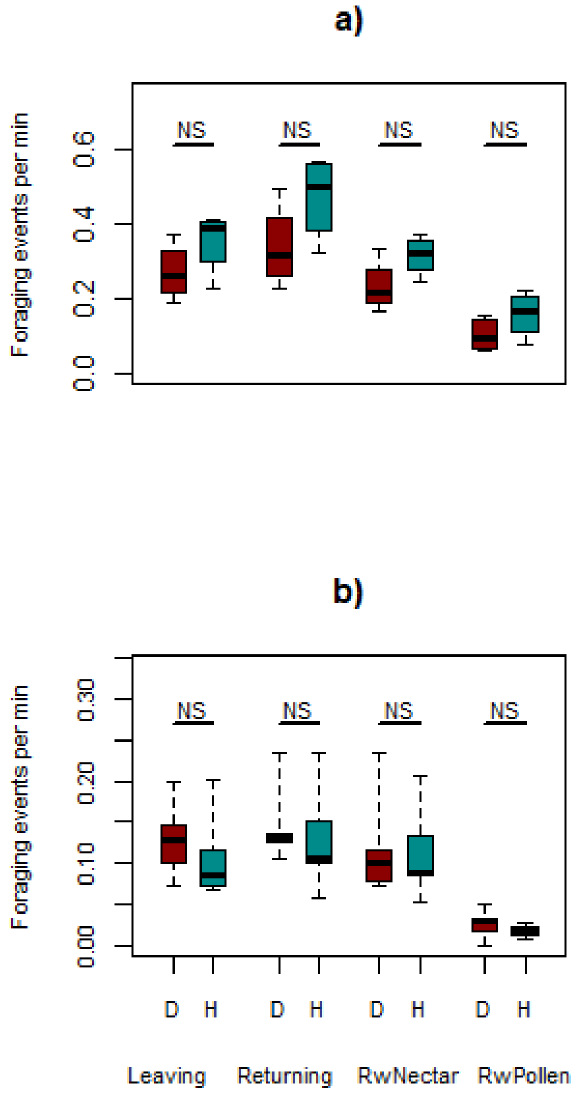


**Figure S2**. The total foraging records in size-diverse and -homogeneous colonies. Shown for Trial 1 (a) and Trial 2 (b). Colony totals shown separately for homogeneous (‘H’) and diverse treatments (‘D’). Foraging records separated into observations in which an individual was: ‘Leaving’ = leaving the nest by darting out, assumed to be on route to forage; ‘Returning’ = returning to the nest (total); ‘RwNectar’ = returning to the nest without pollen, so assumed to be nectar foraging; ‘RwPollen’ = returning to the nest with pollen attached to pollen baskets, demonstrably pollen foraging. Thick black lines = medians; boxes = interquartile ranges; dashed whiskers = ranges. NS = $p > 0.05$, Wilcoxon Rank Sum Test.

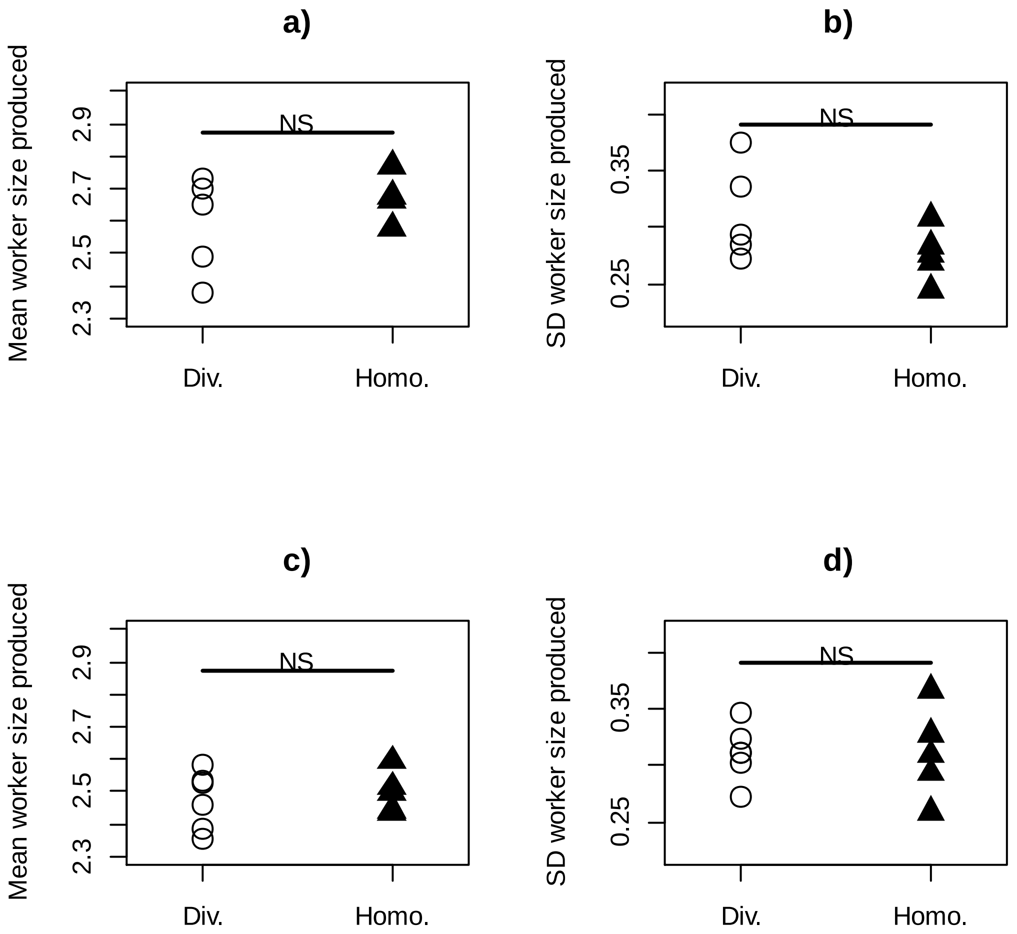


**Figure S3.** The influence of colony body size distribution on the body size of newly emerging bees. Results shown separately for Trial 1 a), b); and Trial 2 c), d). Plots a) and c) show the mean size of worker produced over the course of the experiment per colony, for each treatment. Plots b) and d) show the standard deviation of worker size produced over the course of the experiment per colony, for each treatment. Homo. = size-homogeneous colonies; Div. = size-diverse colonies. NS = p > 0.05, Wilcoxon Rank Sum Test.

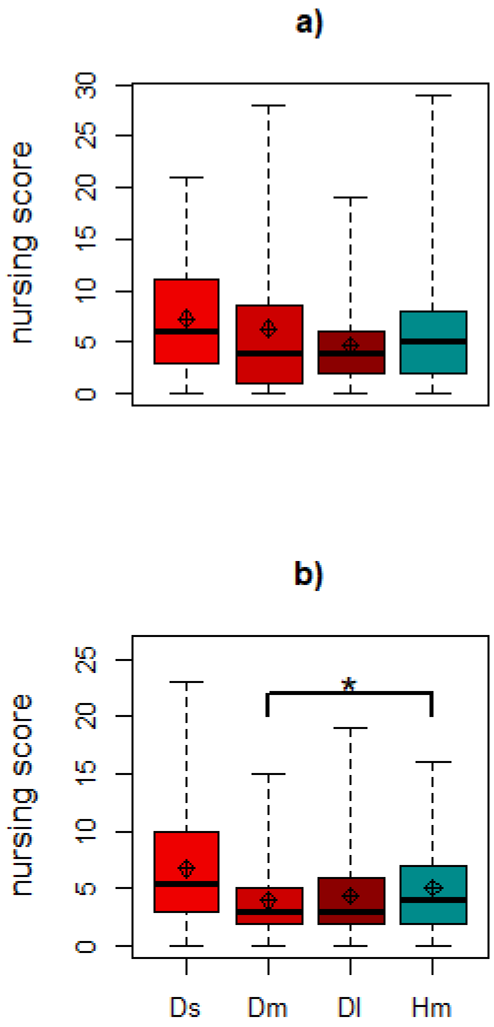


**Figure S4**. Nursing behavior as function of body size and colony composition. Shown for Trial 1 in a) and Trial 2 in b). Nursing records based on frequency of observations when a given worker was tending to larvae or pupae. Ds = small workers from diverse colonies; Dm = medium workers from diverse colonies; Dl = large workers from diverse colonies; Hm = medium workers from homogeneous colonies. Diamonds = means; thick black lines = medians; boxes = interquartile ranges; dashed whiskers = ranges. * $p < 0.05$, Wilcoxon Rank Sum Test.
